# A Circulating TLR2^pos^ CD14^neg^ CD16^neg^ “Unclassified Subset” is Decreased in Multiple Myeloma Patients and May Comprise CD163^pos^ Dendritic Cells

**DOI:** 10.64898/2026.07.21.739900

**Authors:** Mie Wolff Kristensen, Sarah Lindhøj Kvorning, Holger Jon Møller, Marianne Hokland, Thomas Vorup-Jensen, Morten Nørgaard Andersen

**Author notes:** **Corresponding author**: Morten Nørgaard Andersen.

## Abstract

**Background:** Strategies to define human monocytes by flow cytometry vary considerably across studies. Recently, toll-like receptor 2 (TLR2) has been proposed as a marker to identify “all monocytes” in human peripheral blood. However, the TLR2-defined monocytes also contained a previously ignored TLR2^pos^CD14^dim/neg^CD16^neg^ population, which we termed the unclassified subset (UCS).

**Methods:** Peripheral blood mononuclear cells (PBMCs) from healthy donors and patients with multiple myeloma (MM) or monoclonal gammopathy of undetermined significance (MGUS) were analyzed by multiparameter flow cytometry using TLR2^pos^ gating. PBMCs from additional healthy donors were analyzed to characterize the UCS population, including the impact of using either TLR2^pos^ or a negative selection-based gating strategy.

**Results:** The TLR2^pos^ CD14^dim/neg^ CD16^neg^ UCS population was present in healthy controls, MGUS, and MM patients. The UCS expressed the monocyte-macrophage scavenger receptor CD163 and was significantly reduced in MM patients compared to healthy donors (P<0.002). Further phenotypic characterization in healthy blood donors revealed that approximately 80% of UCS cells expressed CD163 at levels comparable to classical monocytes, yet phenotypically resembled CD163^pos^ dendritic cells (DCs). Importantly, gating strategies influenced the composition of the UCS: negative selection-based gating captured all DC subsets, whereas TLR2^pos^ gating primarily included CD1c^pos^ DCs that were highly CD163^pos^.

**Conclusions:** These findings demonstrate that circulating CD163^pos^ CD1c^pos^ DCs are included in the TLR2^pos^ cell population previously described as exclusively monocytes, highlighting the impact of gating strategy on monocyte subset identification. Further, the lower level of TLR2^pos^ CD14^dim/neg^ CD16^neg^ CD163^pos^ cells in MM patients may represent decreased levels of circulating DCs that may contribute to the immune dysregulation in this disease.

## Introduction

Multiple Myeloma (**MM**) is the second most common hematological malignancy, characterized by clonal expansion of malignant plasma cells within the bone marrow. Recent studies have highlighted the critical importance of the bone marrow immune microenvironment in MM pathogenesis, disease progression, and therapeutic resistance.^1, 2^

Among key cellular components shaping this tumor microenvironment are mononuclear phagocytes such as monocytes, macrophages, and dendritic cells (**DCs**), which play important roles in a variety of conditions, including cancer.^3–6^

Tumor-associated macrophages (**TAMs**), particularly those expressing the hemoglobin-haptoglobin scavenger receptor CD163, have been associated with an immunosuppressive tumor environment and adverse outcomes in several cancers, including MM.^7–9^ Furthermore, CD163 expression on circulating monocytes, as well as increased serum levels of soluble CD163 (sCD163), have been associated with poor prognosis in MM and other malignancies.^10–14^ This highlights the clinical and immunological relevance of monocytes and macrophages in MM and in its benign precursor state, monoclonal gammopathy of undetermined significance (MGUS).

In human peripheral blood, monocytes are typically divided into three subsets based on their expression of CD14 (lipopolysaccharide receptor) and CD16 (FcγRIII): classical monocytes (**CM**, CD14^pos^CD16^neg^), intermediate monocytes (**IM**, CD14^pos^CD16^pos^), and non-classical monocytes (**NCM**, CD14^dim^CD16^pos^).^15,16^ CD163 expression is predominantly found on CM and IM, whereas NCM express very low levels.^11,14^ Identification of these subsets by flow cytometry, however, remains methodologically inconsistent across studies and include, e.g., negative selection, CD14^pos^, and light-scatter-based gating strategies. More recently, Toll-like receptor 2 (**TLR2**) has been proposed as a robust pan-monocyte marker that identifies monocytes (TLR2^bright^) while excluding lymphocytes (TLR2^neg^) and neutrophils (TLR2^dim^).^17^

In a recent study examining monocyte subsets in MM, MGUS, and healthy donors,^11^ we observed a previously overlooked TLR2^pos^CD14^dim/neg^CD16^neg^ population within the TLR2^pos^ monocytes, which we termed “unclassified monocytes” (UCM). However, as shown here, our studies revealed that this population contained mainly non-monocytic cells, and thus we renamed it the “unclassified subset” (**UCS**). Because these cells expressed classical myeloid markers such as CD163, but not classical monocyte markers CD16 and CD14, we hypothesized that this population could represent a subset of circulating TLR2^pos^ myeloid DCs.^18–21^

DCs are a heterogeneous family of professional antigen-presenting cells comprising conventional DCs (cDCs) and plasmacytoid DCs (pDCs). Historically, peripheral blood cDCs are further divided into CD1c^pos^ (cDC2) and CD141^pos^ (cDC1) subsets,^22^ but recent studies have shown that CD1c^pos^ DCs can be further divided based on their expression of CD5, CD14, and CD163.^23–25^

In general, studies have indicated compromised DC function in MM, with impaired differentiation and activation, leading to immunosuppression. While studies have shown a general decrease in the total number of circulating DCs in MM patients compared with healthy individuals, knowledge of the specific cDC subsets remains incomplete.^22,26,27^

To our knowledge, the UCS population (TLR2^pos^CD14^dim/neg^CD16^neg^CD163^pos^) has not previously been investigated. Thus, the present study aimed to further characterize the UCS in human peripheral blood mononuclear cells, evaluate whether it represented circulating DCs, and assess how different flow cytometric gating strategies influence the UCS composition. We found that the TLR2^pos^ UCS was significantly reduced in MM compared to healthy donors, and subsequent analysis in healthy donor samples revealed that this subset mainly consisted of non-monocyte CD163^pos^ CD1c^pos^ DC-like cells. Further studies are warranted to understand the biological and clinical significance of TLR2^pos^ UCS in MM.

## Methods and Materials

### Patient and healthy donor samples

Flow cytometry data on peripheral blood mononuclear cells (PBMCs) from patients with MM, MGUS, and healthy controls were collected as described previously.^11^ Briefly, the study included patients with multiple myeloma (n = 32) and MGUS (n = 8) from the Department of Hematology, Aarhus University Hospital (AUH), Aarhus, Denmark. Samples from healthy donors (n = 16) were included from the blood bank, Department of Clinical Immunology, AUH, Denmark. PBMCs were isolated and stored at −80°C until analysis. The study was approved by the local ethics committee of the Central Denmark Region (ref. 1-10-72-144-17). The flow cytometry data obtained in the previous study were reanalyzed for the present study with special focus on the UCS.

Further, PBMCs were also isolated from anonymized healthy blood donor buffy coats obtained from the blood bank, Department of Clinical Immunology, AUH, Aarhus, Denmark (project no. 77) in accordance with Danish law.

### Isolation of PBMCs

The healthy donor PBMCs were isolated as previously described.^28^ Briefly, cells were isolated using density gradient centrifugation on a Histopaque-1077 gradient (Sigma-Aldrich, Munich, Germany) according to the manufacturer’s instructions. After isolation, cells were washed twice in PBS with 2% fetal calf serum (v/v, FCS, Lonza, Basel, Switzerland) and stored at −150°C in RPMI-1640 medium with penicillin/streptomycin (100U/100μg pr. mL, Lonza) containing 20% FCS and 10% DMSO (v/v) (Sigma Aldrich). Cells were thawed at 37°C, washed once in PBS with 20% FCS, and resuspended in stain buffer (PBS with 0.5% bovine serum albumin (w/v) (BSA, Calbiochem, San Diego, CA) and 0.09% NaN_3_ (v/v) (Sigma Aldrich) at a final cell concentration of 10×10^6^/mL.

### Staining of PBMCs and flow cytometry

Blocking reagents were added to all samples: Fc receptor block (purified human IgG, Beriglobin, CSL Behring GmbH, Marburg, Germany) was applied at 100µg/ml.^28^ For blocking of cyanine-tandem fluorophore non-specific binding, “Oligo-Block” (phosphorothioate DNA oligo: 5’-GGGGGAGCATGCTGGGGGGG-3’, Sigma-Aldrich) was used as previously described,^29^ and incubated for 15min at 4°C. After blocking, primary antibodies were added (see **Supplementary Table 1**) without removing the blocking reagents. Samples were incubated in the dark for 30min at 4°C, in a final staining volume of 100µl. After incubation, samples were washed in stain buffer, fixed with 0.9% (v/v) formaldehyde in PBS for 15min, washed again, and resuspended in PBS for analysis.

As compensation controls, we used single-stained OneComp eBeads (eBiosciences, San Diego, CA) with positive and negative beads for all primary antibodies. 7-AAD compensation was done using 100% methanol fixed cells with a mixed positive and negative/unstained population. Samples were analyzed on an LSRFortessa flow cytometer (BD Biosciences, San Jose, CA). All fluorescence values were recorded as voltage pulse area, and all events were within the linear range of the detectors. All fluorescence signal intensities are reported as median fluorescence intensity (MFI). A minimum of 500,000 events were acquired for all PBMC samples and 30,000 events for compensation samples. Data was compensated and analyzed, and figures were made using FlowJo^TM^ 10.8.1 for Mac (FlowJo, LLC, Ashland, OR). Figure layout was done in Affinity Designer (Affinity Serif, Nottingham, UK).

The negative selection gating strategy for identification of DC subsets in samples from healthy donors is shown in **Figure 2**, and the TLR2^pos^ gating and negative selection gating strategies for identification of monocytes are shown in **Supplementary Figure 2**.

Due to the very low frequency of DCs in peripheral blood, the frequency of these subsets in flow cytometry data is reported here as the number of cells per 10,000 CD45-positive live single cells.

### Statistics

All statistical analyses were performed using Prism 8^TM^ for Mac (GraphPad Software, San Diego, CA). Normal distribution was evaluated using QQ-plots and the Shapiro-Wilk Test. Data is shown as mean ± 95% CI. For comparison, we used mixed-effect ANOVA with Tukey’s multiple comparisons post-test and paired analysis. Appropriate multiple comparison corrections were applied to all relevant data. A P-value <0.05 was considered statistically significant, and only significant p-values are shown in the figures.

## Results

### “Unclassified Monocytes” in Multiple Myeloma Patients

In our previous report, focused on CD163 expression on monocytes in patients with MGUS or MM,^11^ we defined the “total monocyte population” as TLR2^pos^ with subsequent identification of the three conventional monocyte subsets based on CD14 and CD16 expression. No significant differences were found in the frequencies of classical, intermediate, and non-classical monocytes between healthy donors, MGUS, and MM patients.

In the present study, we re-analyzed the flow cytometry data and observed a population of CD14^dim/neg^ CD16^neg^ TLR2^pos^ cells, which we named the “unclassified monocytes” (UCM, **Figure 1A**). Interestingly, in contrast to the three conventional monocyte subsets, the UCM subset frequency was significantly lower in newly diagnosed MM patients, 2.72% (1.65; 3.80), compared to healthy controls, 4.6% (3.9; 5.2) (P=0.01). Similar results were found for MM patients in disease remission, and at disease relapse (P<0.001, **Figure 1B**). Further, the frequency of CD163^pos^ UCM was lower in newly diagnosed MM patients, 75.0% (66.1; 84.0), compared to healthy controls, 87.7% (85.2; 90.1) (P=0.02). Otherwise, there were no differences between groups in the frequency of CD163^pos^ cells (**Figure 1C**) or the expression levels (**Figure 1D**). Based on these results, we wanted to further investigate and characterize the poorly described population of UCM. Since additional samples from the previous study^11^ were not available, we decided to further characterize the UCM population using healthy blood donor PBMCs.

**Figure 1:**
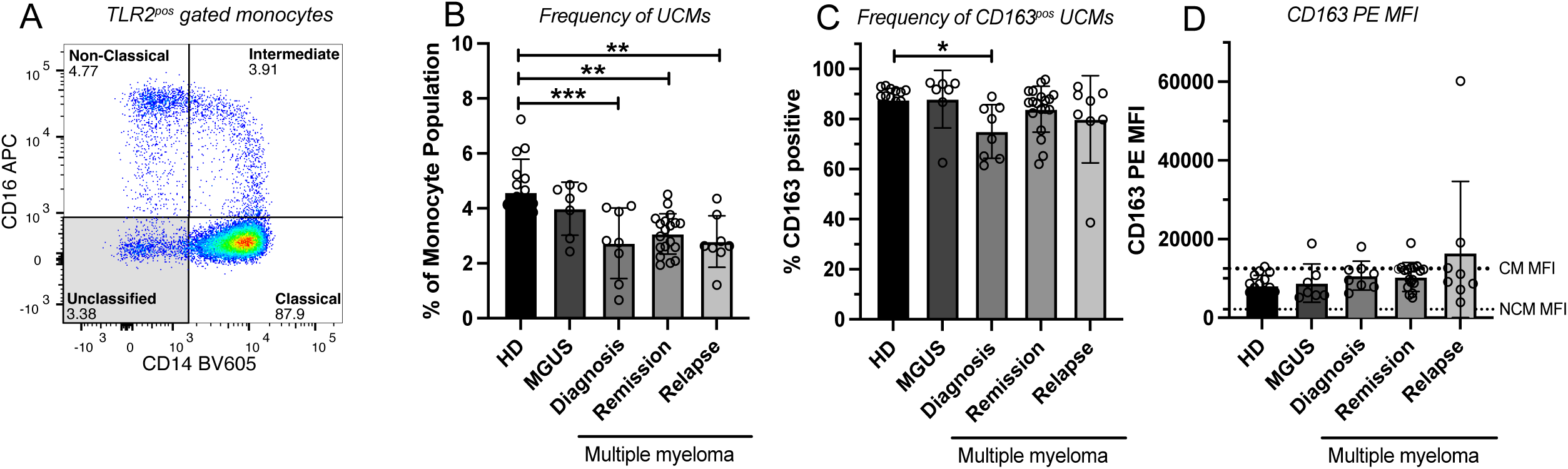
“Unclassified Monocytes” and CD163 expression in MGUS and Multiple Myeloma Results on “Unclassified monocytes” (UCM, TLR2^pos^ CD14^dim/neg^ CD16^neg^) from Healthy Donors (N=16), and patients with monoclonal gammopathy of unknown significance (MGUS, N=7), or multiple myeloma using data from Kvorning et al (2018).^5^ Multiple myeloma patients were categorized as: at diagnosis (N=8), in remission (N=19), or relapsed (N=8). A) TLR2^pos^ gated monocytes: A representative example of the TLR2^pos^ gated monocytes in a sample from a patient with newly diagnosed multiple myeloma. UCM were defined as _CD14_dim/neg _CD16neg_. B) UCM counts as a percent of the TLR2^pos^ defined “all monocytes” population. C) Percent CD163-positive UCM of all UCM. D) CD163 expression level (PE MFI) of the CD163^pos^ subset of UCM. The median CD163 expression levels of Classical (CM MFI) and Non-classical (NCM MFI) monocytes are marked with dotted lines. In Kvorning et al (2018), no difference in CD163 expression level of CM, IM, and NCM between patient categories was found. Data is shown as mean and standard deviation. P-values are only shown for statistically significant results. The used gating strategy can be found in Supplementary Figure 2 in the original paper.^5^ CD163^pos^ cells were gated based on a fluorescence minus one (FMO) control.

These data on healthy donor PBMCs are presented below and showed that the UCM population likely consisted of non-monocytic cells, and therefore, this population will be named the “unclassified subset” (**UCS**) from here on.

### Characterization of the “Unclassified Subset” in online available data

Previously, Mair et al. (2018)^30^ published an optimized multicolor immunofluorescence panel (OMIP) for characterization of circulating DCs in healthy donors, and they included CD163 as a marker. Using their dataset (available in FlowRepository, ID: FR-FCM-ZYC2), we were not able to directly identify our UCS population, since the used panel did not include TLR2. However, our analysis of the Mair data did show that a large fraction of CD1c^pos^ DCs were CD14^dim/neg^ CD16^neg^ and CD163^pos^, thus indicating that CD1c^pos^ DCs could be included in our UCS (see **Supplementary Figure 1**). Expression levels of additional markers such as CD80, CD86, CD40, and CCR7 were also similar between CD1c^pos^ DCs and a negative selection-gated UCS.

### The “Unclassified Subset” largely consists of Dendritic Cells

Since Mair et al. (2018) did not include TLR2 in their panel, a direct comparison of the DC and UCS populations of interest was not possible using their data. Thus, we created our own flow cytometry panel based on the panel from Mair et al. (2018)^30^ to investigate whether the UCS (TLR2^pos^ CD14^dim/neg^ CD16^neg^) was comprised of DCs. Using this panel, we identified the DC subsets described by Mair et al.^30^ in healthy donor PBMC samples (n = 10): pDCs, CD141^pos^ DCs, CD1c^pos^ DCs, and double-negative (DN) DCs (**Figure 2A**). Of the four subsets, only the CD1c^pos^ showed expression of CD163, and TLR2 was only expressed by CD1c^pos^ and DN DCs (**Figure 2B**). When displaying the four DC subsets on a CD14-CD16 plot, it was clear that the majority of DCs (except for DN DCs) were CD14^dim/neg^ CD16^neg^, similar to the “Unclassified Subset” (**Figure 2C**).

**Figure 2:**
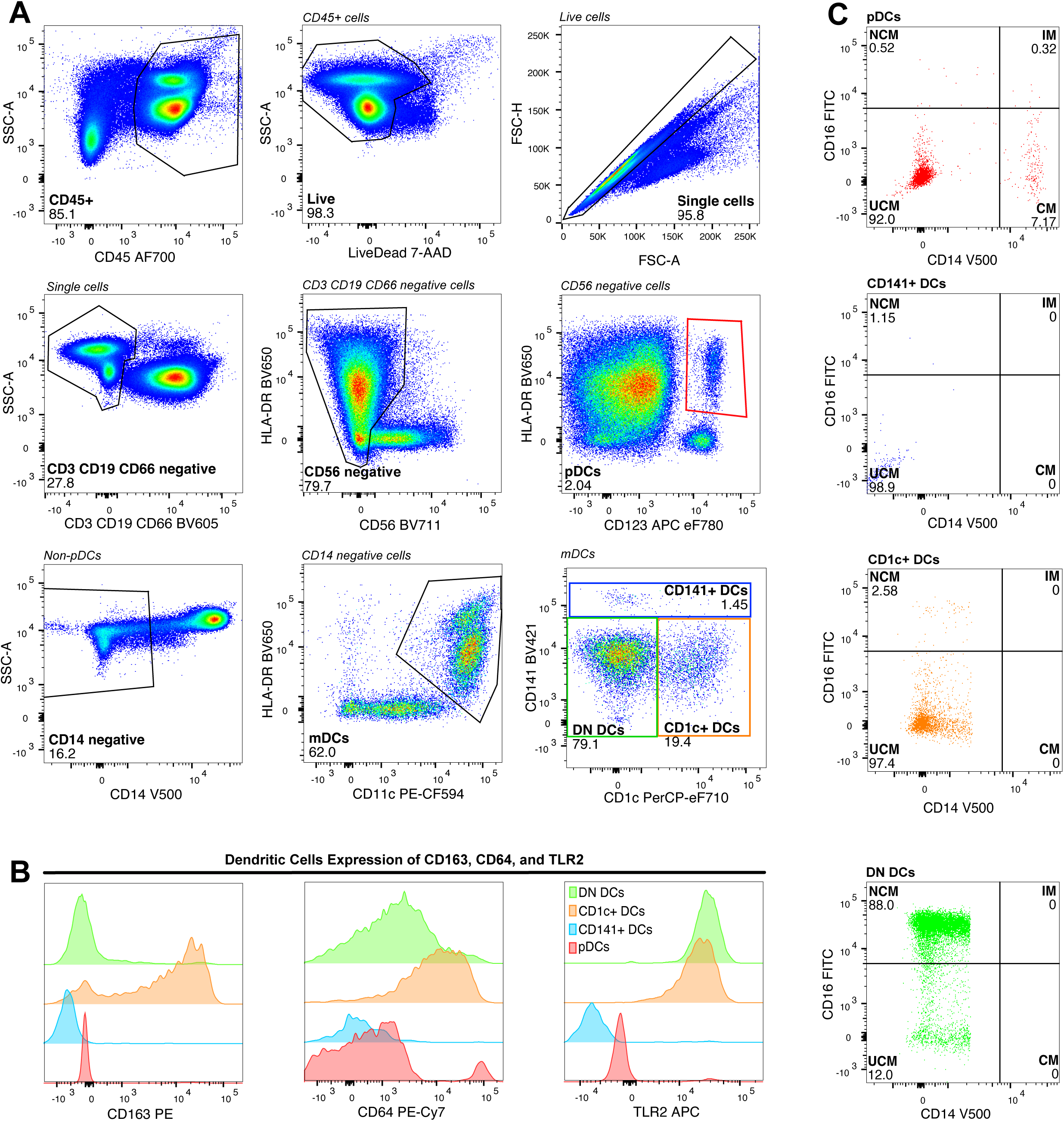
Gating Strategy for Identification of Dendritic Cell Subsets Gating Strategy for identification of DC subsets in samples from healthy donors (N=10), representative example. A) The used gating strategy based on the OMIP by Mair et al (2018).^23^ Leukocytes are defined as live CD45^pos^ single cells. Dendritic cells are defined as lineage negative (Lin^neg^: CD3 CD19 CD66 CD56 CD14 negative) and subsets are defined as plasmacytoid DCs (pDCs, Lin^neg^ HLA-DR^pos^ CD123^pos^) and myeloid DCs (mDCs, Lin^neg^ HLA-DR^pos^ CD11c^pos^). mDCs are further divided based on their CD141 and CD1c expression into CD141^pos^ DCs, CD1c^pos^ DCs, and double negative (DN) DCs. B) Expression of CD163, CD64, and TLR2 for all four DC subsets. C) Distribution of the four DC subsets in a monocyte plot (CD14 vs CD16). Gating strategies for TLR2^pos^ and negative selection defined monocytes are shown in Supplementary Figure 2.

Analysis of the four DC subset frequencies (displayed as cell count per 10,000 CD45^pos^ Live single cells) showed similar results for pDCs and CD1c^pos^ DCs at 56.6 (37.4; 75.8) and 46.0 (27.0; 64.9), respectively. DN DCs were more abundant at 150 (102; 199), while CD141^pos^ DCs were the least frequent at 4.5 (2.6; 6.4) (**Figure 3A**).

**Figure 3:**
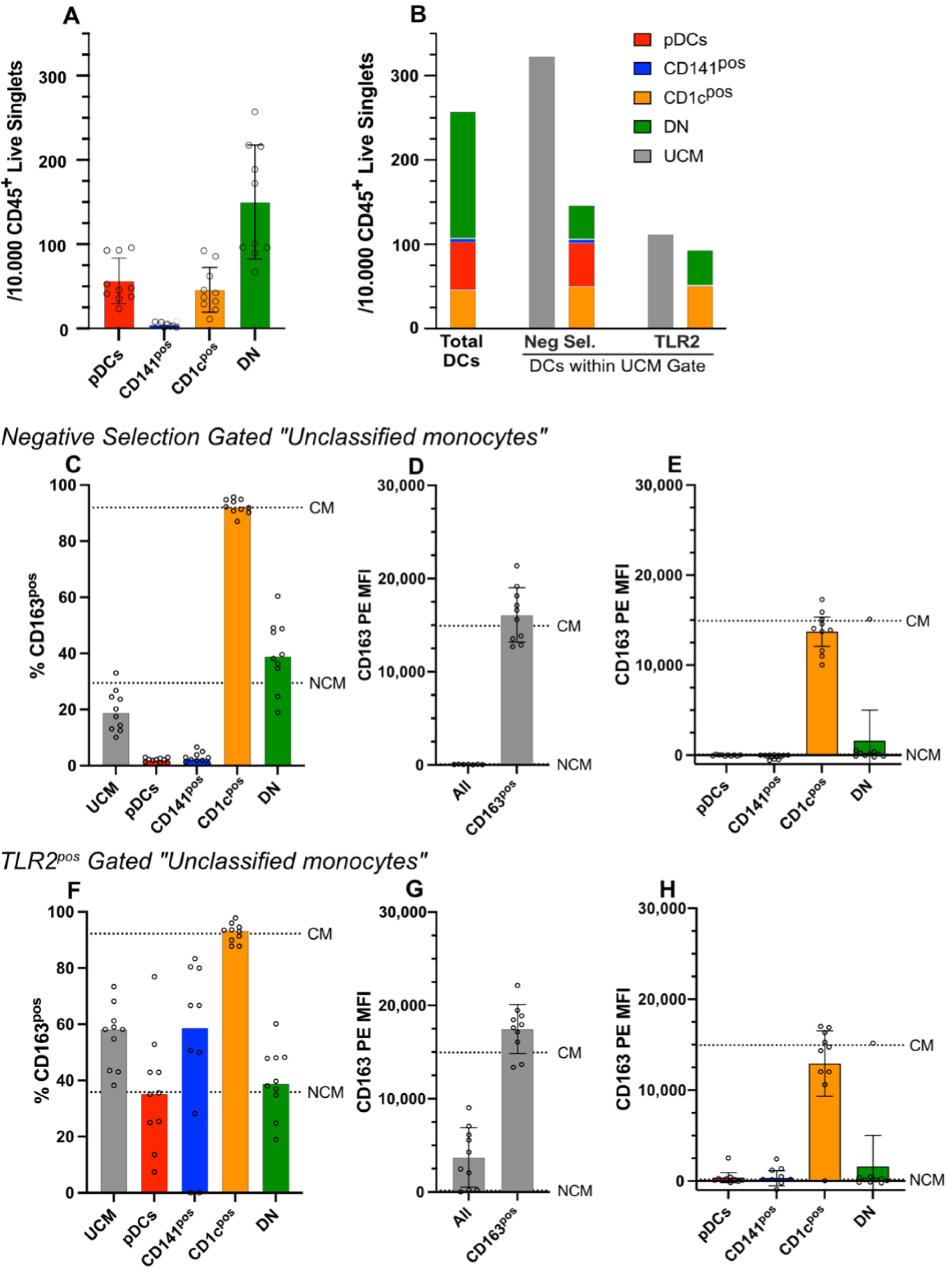
Dendritic Cells and the “Unclassified Subset” A) Frequency of dendritic cells (DCs) in healthy donors (N=10) shown as no. of cells per 10,000 CD45^pos^ live single cells. The DCs were gated as shown in Figure 2. B) The first column shows the added frequencies of total DCs shown in A). The following columns show the frequency of DCs included in the “unclassified subset” (UCS) defined by either negative selection (neg. sel.) or TLR2^pos^ gating. The frequencies of UCS for both populations are shown as a reference (grey column). Both gating strategies excluded DN DCs, while TLR2^pos^-based gating excluded almost all CD141^pos^ DCs and pDCs. For negative selection gating, non-DCs included in the UCS gate were CD123^pos^ HLA-DR^neg^ cells, most likely basophils and eosinophils, and a population of unknown cells (CD11c^neg^ HLA-DR^neg^ Lin^neg^). For TLR2^pos^-based gating, non-DCs included in the UCS gate were a small number of T cells, NK cells, B cells, and granulocytes, as well as a small population of unknown cells (CD11c^neg^ HLA-DR^neg^ Lin^neg^). For representative examples of UCS gated by neg. sel. and TLR2-based gating see Supplementary Figure 2. C-H) Results are shown for both UCS defined by negative selection (C-E), and UCS defined by TLR2-based gating (F-H): C+F) The frequency of CD163^pos^ cells among UCS and the pDCs, CD141^pos^, CD1c^pos^, and DN DCs included in this population. D+G) CD163 expression levels as CD163 PE MFI for the entire UCS population, and for the CD163^pos^ UCS population. E+H) CD163 expression levels as CD163 PE MFI for the DC subsets included among the UCS population. In C-H) Mean values for frequency or CD163^pos^ MFI of non-classical monocytes (NCM) and classical monocytes (CM) are shown as dotted lines. Data is shown as mean with standard deviation for healthy donors (N=10). P-values are only shown for statistically significant results.

As described above, we wanted to investigate whether the UCS consisted of DCs. However, in the literature, gating strategies for defining human monocytes vary greatly, but negative selection (exclusion of lineage marker-positive cells) has been considered the gold standard by several researchers.^17,31^ Thus, to thoroughly assess the composition of the UCS, we choose to analyze our flow cytometry data using both a TLR2^pos^ and a negative selection-based gating strategy to define the total monocyte population.

When “all monocytes” were defined using a *negative selection gating strategy*, the UCS population contained 45.2% (33.3; 57.0) DCs (**Figure 3B**), and roughly all pDCs, CD141^pos^, and CD1c^pos^ DCs were included in the UCS population. However, 71.6% (64.2; 79.0) of DN DCs were excluded.

When “all monocytes” were defined by *TLR2^pos^ gating*, the overall UCS frequency was lower compared to negative selection gating but contained a higher proportion of DCs: (83.4% (75.3; 91.5). In this approach, approximately 99% of pDCs and CD141^pos^ DCs, and 83.7% (79.3; 88.1) DN DCs were excluded, while no CD1c^pos^ DCs were excluded and comprised 50.9% (15.2; 93.5) of the UCS (Figure 3B).

These results are further illustrated in **Supplementary Figure 2**, where the DC gating strategy was applied to the UCS population defined by either the negative selection or TLR2^pos^ monocyte gating. Here, expression levels of HLA-DR, CD11c, CD64, and CD163 were similar between CD1c^pos^ DCs and the TLR2^pos^ gated UCS (Supplementary Figure 2D-J).

Overall, most DCs were CD14^dim/neg^ CD16^neg^ and thus located in the UCS gate on a monocyte plot, except the DN DCs, as also shown in Figure 2C.

Further, the negative-selection gating UCS included all pDCs, CD1c^pos^ DCs, and CD141^pos^ DCs. However, only ∼45% of cells were DCs, and thus the population contained >50% of other cell types. In contrast, TLR2^pos^ gated UCS comprised ∼85% DCs, but virtually all CD141^pos^ and pDCs were excluded. Thus, these results emphasize that the markers and gating strategy chosen to define monocyte subsets can significantly impact which DCs are included.

### CD163 Expression by the “Unclassified Subset” and Dendritic Cell Subsets

As shown above, for both controls and patients with MGUS and MM, the majority of cells in the UCS population were CD163^pos^. Since CD163 expression in the tumor microenvironment and on circulating monocytes has been associated with prognosis in several cancers, we further investigated CD163 expression on UCS and the four DC subsets in healthy donors. Applying the two investigated gating strategies, the frequency of CD163^pos^ UCS was lower for negative selection monocyte gating (19.1% (14.1; 24.2), **Figure 3C**) compared to TLR2^pos^ gating (56.1% (47.4; 64.8), **Figure 3F**).

CD163 expression was analyzed for UCS, including both the whole-population MFI and the CD163^pos^ population MFI. For the negative-selection-gated total UCS population, CD163 expression was very low and similar to that of NCM, while CD163 expression of the CD163^pos^ UCS population was similar to that of CM. A similar pattern was seen for TLR2^pos^-gated UCS, but with somewhat higher MFI levels (**Figure 3D+G**).

For the DC1 subsets, CD1c^pos^ DCs had the highest CD163^pos^ fraction (>90%) and CD163 MFI level for both gating strategies, similar to that of CM. The expression of CD163 and the frequency of CD163^pos^ pDCs, CD141^pos^, and DN DCs were lower or similar to that of NCM. The frequency of CD163^pos^ DCs was higher within TLR2^pos^-gated monocytes compared to negative selection gating (**Figure 3C, E, F, and H**).

Taken together, using TLR2 to define “all monocytes” resulted in inclusion of cells with higher CD163 expression, both percent positive and MFI, compared to negative selection.

### t-SNE analysis of DCs and Monocyte subsets

To further characterize the monocyte and DC subsets, including their CD163 expression, we used t-SNE analysis to objectively evaluate the expression of the panel of surface markers (**Figure 4**). The populations were defined based on gating strategies shown in **Figure 2, Supplementary Figure 2,** and **Supplementary Figure 3**, overlaid on the t-SNE plot (**Figure 4A**) and used to generate heatmaps showing expression patterns of the selected markers (**Figure 4B-J**).

**Figure 4:** t-SNE analysis of Dendritic Cell subsets. A) t-SNE plot with all CD45^pos^ live cells from a representative healthy donor using FlowJo. Monocyte and DC subsets were defined as shown in Figure 2 and Supplementary Figure 2. Remaining subsets were defined as shown in **Supplementary** Figure 3. Monocyte and DC subsets were overlaid on the t-SNE plot to highlight co-localization among subsets. pDCs, CD1c^pos^, CD141^pos^, and DN DC populations are marked with colored rings. * Double negative (DN) DCs co-localized with the non-classical monocyte population and cannot be seen. † Not all “unclassified” cells can be seen clearly due to co-localization/overlay with the four DC subsets. B-J) t-SNE heat maps showing expression levels of CD14, CD16, CD11c, HLA-DR, TLR2, CD163, CD123, CD1c, and CD141.

The CM, IM, and NCM clustered nicely together, while the UCS were more spread out and clustered with both DCs and monocyte subsets, thus supporting the finding that UCS consist of several different cell types for the negative selection gating strategy. However, the majority of UCS co-localized with DC subsets.

The heatmap for TLR2 (**Figure 4F**) showed expression co-localized with monocyte subsets and CD1c^pos^ DCs (but not with other cell types). The heatmap for CD163 (**Figure 4G**) showed expression co-localized with CM and IM, and with CD1c^pos^ DCs (but not with other cell types). Thus, this analysis supported the results from our manually gated analyses described above. Taken together, these data support the conclusion that TLR2^pos^ UCS mainly comprises CD1c^pos^ CD163^pos^ DCs, also known as cDC3 in the newer nomenclature.

## Discussion

Previous studies have demonstrated immune dysfunction in multiple myeloma patients, including impaired DC proliferation, migration, and maturation, which may contribute to the reduced anti-tumor immunity and, at least in part, explain the limited clinical efficacy of DC-based vaccines.^22,32^ In the present study, we analyzed blood samples from healthy donors and from patients with MGUS or MM to characterize a previously ignored subpopulation within the TLR2^pos^-defined monocytes. These CD14^dim/neg^CD16^neg^ cells were initially termed the “unclassified monocyte subset.”

Notably, this subset was significantly decreased in MM patients compared with healthy donors, in contrast to the three conventional monocyte subsets, whose frequencies remained unchanged between groups, as previously shown.^11^ As described here, further analyses in healthy donor samples revealed that this TLR2^pos^ “unclassified” population included mostly dendritic cells, particularly CD163^pos^ DCs. Based on this, the population was redefined as the “unclassified subset” (**UCS**), and we aimed to further characterize this previously ignored population, including the impact of different strategies for gating of monocytes by flow cytometry.

Our results demonstrated that when “all monocytes” were defined as TLR2^pos^, based on Shirk et al.^17^, the UCS predominantly contained CD163^pos^ DCs (CD163^pos^ CD1c^pos^ CD14^dim/neg^ cDCs or cDC3s).^23–25,33^ In contrast, when defining “all monocytes” by a negative selection strategy, the UCS contained several non-monocytic subsets, including DCs with lower CD163 expression levels. These findings emphasize that the gating strategy used to define “all monocytes” determines which non-monocytic cells are included in the UCS population.

Based on the characterization of the UCS in healthy donor samples, we hypothesize that the reduced UCS population observed in MM patients could reflect a decrease in a subset of circulating CD163^pos^ cDC3s. This has not been reported previously, and further studies are needed to understand the significance of these cells in MM patients, including their associations with immune dysfunction.

Historically, DCs have been classified based on their surface marker expression,^34–36^ but recent high-dimensional analyses, including single-cell sequencing, have identified additional subsets within the CD1c^pos^ DCs.^23–25,37^ They suggest that CD1c^pos^ DCs are two distinct subsets: CD163^neg^ cDC2 and CD163^pos^ cDC3.^23,24^ Because the circulating CD1c^pos^ DCs identified in the present study were mostly CD163^pos^, they most likely belong to the cDC3 subsets.

However, while our data suggest that the decreased UCS frequencies in MM may reflect a reduction in circulating cDC3s, this hypothesis requires confirmation in MM patient samples using expanded flow cytometry panels and functional assays assessing, e.g., T-cell stimulatory capacity. It would also be informative to compare circulating and bone marrow-resident DCs to better understand compartment-specific alterations in MM patients. Further, a limitation of the present study is the use of a relatively small flow cytometry panel, adapted from Mair et al. 2018,^30^ which was feasible using an LSRFortessa cytometer. More recently, larger panels have been described for use on more advanced systems.^23–25,38^ These studies have identified additional discriminative markers, such as CD88 and CD89, which should be included in future work to more precisely differentiate cDC3 and monocyte populations.^33^ Lastly, the lack of available MGUS and MM samples is a limitation that necessitated the use of samples from healthy blood donors to characterize the UCS population.

CD163 is an established macrophage and DC marker with biomarker and therapeutic relevance, where soluble CD163 has shown potential as a biomarker in MM,^12^ and membrane-bound CD163 may serve as a target for drug delivery. ^39^ However, few studies have focused on monocytes and dendritic cells in MGUS and MM, including their CD163 expression. Studies have highlighted the potential of CD163^pos^ cCD3 in stimulating T cell proliferation and their potential importance in cancer vaccines,^40–42^ yet their role in infection and cancer remains largely unexplored.^21,33^ Our findings of reduced circulating CD163^pos^ UCS/cDC3s fit with existing literature of reduced peripheral blood DCs in MM, and may provide a potential mechanistic link to the impaired DC-mediated immunity described in MM.^26,27^

Despite initial promise, clinical trials for DC-based vaccines have failed in MM; however, most studies used monocyte-derived DCs rather than circulating DC subsets.^41,43^ While data on the significance of circulating DCs in MM is limited, emerging evidence from solid tumor malignancies highlights their anti-cancer potential. For example, low levels of CD1^pos^ DCs were associated with worse outcomes in lung adenocarcinoma,^44^ and intratumoral administration of CD1c^pos^ and CD141^pos^ DCs enhanced antitumor response in melanoma patients who previously progressed on immune therapy.^43^

Murine MM models have shown that combining DC-based vaccines with immunomodulatory imid drugs (IMiDs), dexamethasone, and either anti-PD1 or anti-PDL1 may augment T cell activation.^45,46^ Given the improved T cell activation observed in the triple combination arm, it could be speculated that treatment with IMiDs or dexamethasone would impact circulating DC counts or activation state in MM patients treated with these drugs. In our study, the data did not show any differences for the UCs between newly diagnosed MM (treatment naïve) and relapsed MM or MM patients in remission (treated patients). Thus, our data did not support such a hypothesis, but further studies on larger MM cohorts with dedicated flow cytometry panels would be needed to investigate this aspect further.

The present study also provides a methodological contribution by demonstrating that the flow cytometric definition of “all monocytes” markedly affects the inclusion of DC subsets. Using negative selection gating captured nearly all DC populations (pDCs, CD1c^pos^, CD141^pos^, and a fraction of DN DCs) in the UCS population. However, it also included non-DCs such as eosinophils/basophils (CD123^pos^ HLA-DR^neg^) and some unknown CD11c^neg^ HLA-DR^neg^ cells. In contrast, the TLR2^pos^ UCS included CD1c^pos^ and a subset of DN DCs while excluding pDCs and CD141^pos^ and minor contamination by non-myeloid cells. In general, the TLR2^pos^ UCS was more CD163^pos^ compared to the negative-selection-gated UCS, consistent with greater myeloid specificity. These findings highlight that awareness is needed when designing flow cytometry panels, and that there is need for standardized gating strategies to enable comparability between studies. Here, TLR2^pos^ gating of human blood samples may allow for easy identification of conventional monocyte subsets and a major subset of CD1c^pos^ CD163^pos^ DCs in the UCS population.

**In conclusion**, we have characterized a previously ignored TLR2^pos^ CD14^dim/neg^ CD16^neg^ population in PBMCs from healthy donors and patients with MGUS and MM. This “unclassified subset” exhibited CD163 expression similar to that of classical monocytes and appeared decreased in MM patients. Our further analyses in healthy blood donors indicate that the UCS predominantly contained myeloid CD163^pos^ CD1c^pos^ DCs, most consistent with the cDC3 subset. These findings highlight the importance of choice of gating strategy in defining human monocytes, and further suggest that loss of CD163^pos^ cDC3s could contribute to immune dysregulation in MM patients, which should be verified in future studies.

## Acknowledgment

All experiments were performed using the LSRFortessa flow cytometer (BD Biosciences) at the FACS Core Facility, Aarhus University, Denmark.

## Author Contribution

MNA: conceived the study. MWK and MNA: Designed the study. MWK and SLK performed the Flow Cytometry Experiments. MWK, SLK, MH, HJM, TVJ, and MNA: analyzed and interpreted data. MWK and MNA: drafted the manuscript. All authors provided critical review and approved the final version of the manuscript.

