## Supplementary Materials for "A Circulating TLR2^pos^ CD14^neg^ CD16^neg^ “Unclassified Subset” is Decreased in Multiple Myeloma Patients and May Comprise CD163^pos^ Dendritic Cells"

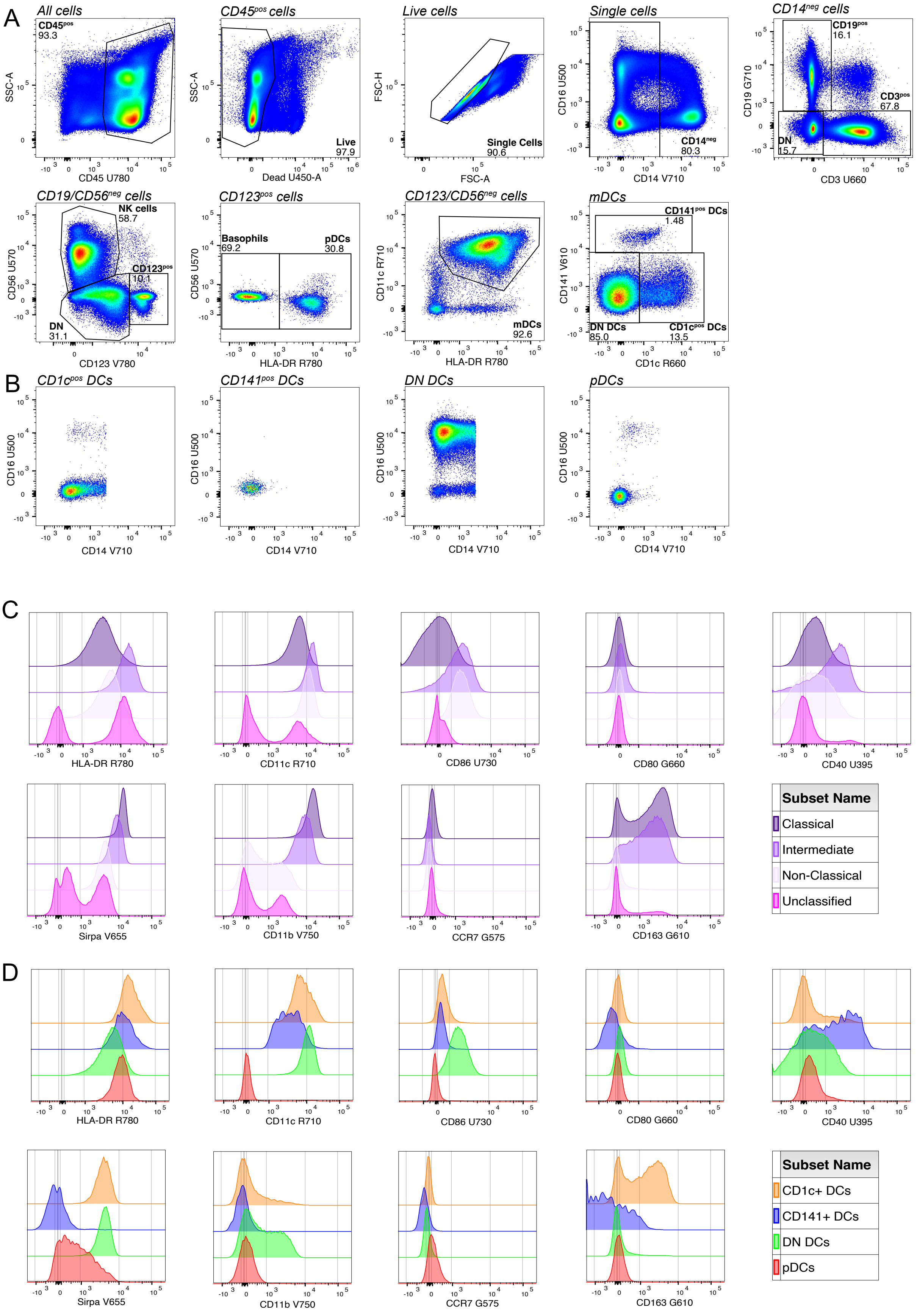


**Supplementary Figure 1**: Identification of CD163$^{pos}$ Dendritic Cells Using Flow Cytometry Data from Mair et al (2018) [22].

A) Gating Strategy for identifying dendritic cell (DC) subsets, which is identical to the one used in the original article. mDCs were defined as lineage negative (CD14$^{neg}$, CD19$^{neg}$, CD56$^{neg}$, CD3$^{neg}$ cells) and mDCs were divided into CD1c$^{pos}$ CD141$^{pos}$, and double negative (DN) DCs. Monocytes were defined as lineage negative (CD19$^{neg}$, CD56$^{neg}$, CD3$^{neg}$ cells.

B) Placement of the four DC subsets in a CD14 versus CD16 plot (monocyte plot)

C) Expression of HLA-DR, CD11c, CD86, CD80, CD40, Sirpa, CD11b, CCR7, and CD163 on monocytes gated by negative selection gating (CD19$^{neg}$, CD56$^{neg}$, CD3$^{neg}$)

D) Expression of HLA-DR, CD11c, CD86, CD80, CD40, Sirpa, CD11b, CCR7, and CD163 on DC subsets.

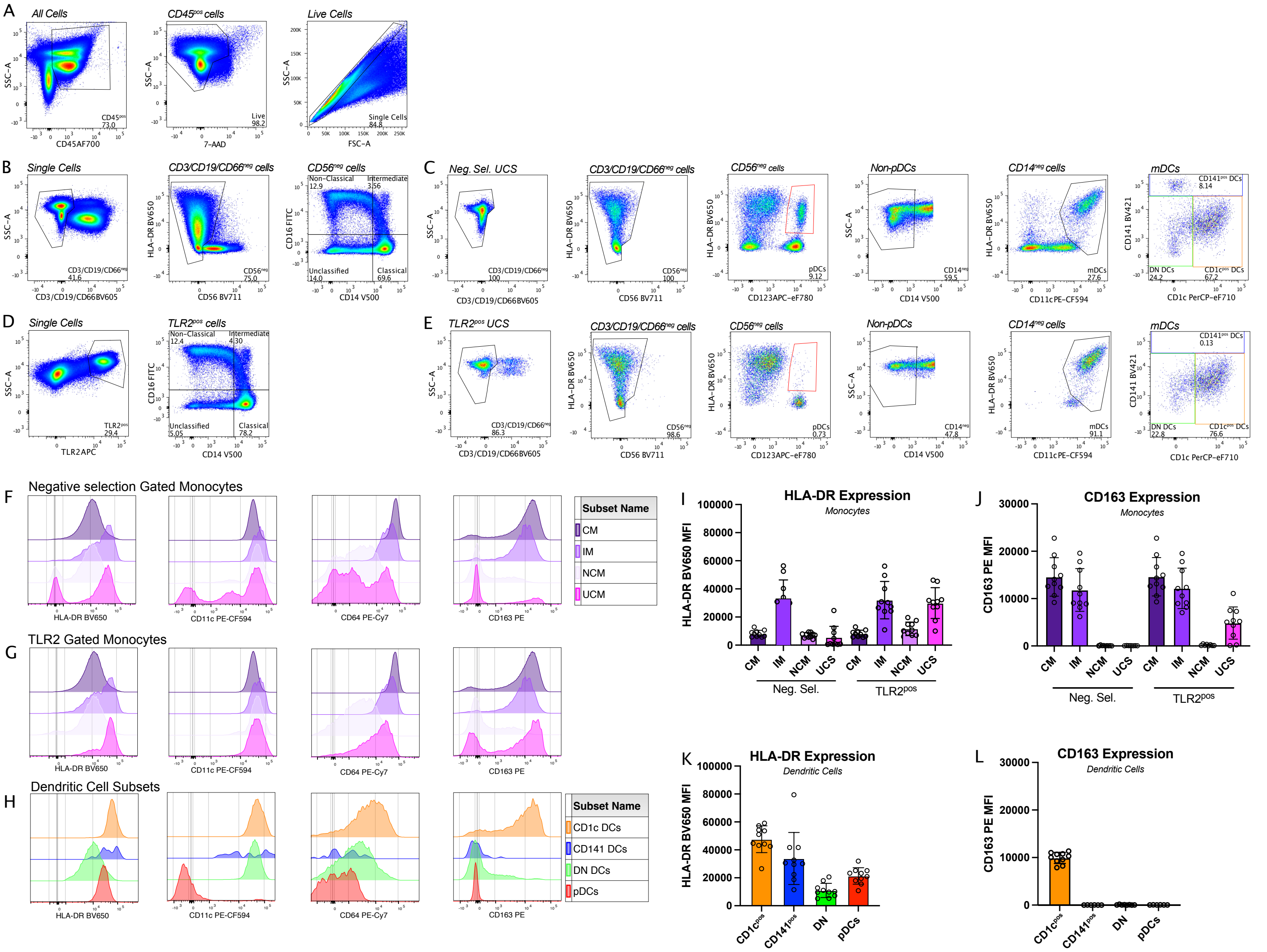


**Supplementary Figure 2**: Dendritic Cells (DCs) within the "Unclassified Subset" (UCS) populations defined by either TLR2$^{pos}$ or negative selection gating
A) General initial gating strategy applied to all samples.
B) Negative selection gating of all monocytes and UCS
C) DC gating applied to the negative selection defined UCS population. It is seen that all DC subsets are included in the UCS population. Non-DCs included in the gate are a CD123$^{pos}$ HLA-DR$^{neg}$ population (~32%), which may comprise basophils and eosinophils, and an unknown CD11c$^{neg}$ and HLA-DR$^{neg}$ Lin$^{neg}$ population.
D) TLR2 gating of all monocytes and UCS.
E) DC gating applied to the TLR2$^{pos}$ defined UCS population. It is that almost all pDCs and CD141$^{pos}$ DCs are excluded, while both CD1c$^{pos}$ and DN DCs are present in the UCS population. Non-DCs included are T (~2%), NK cells (~3%), B cells (~4%), and granulocytes (~2%). They also had a small CD11c$^{neg}$ and HLA-DR$^{neg}$ Lin$^{neg}$ population.
F) Expression of HLA-DR, CD11c, CD16, and CD64 on negative selection gated monocyte subsets
G) Expression of HLA-DR, CD11c, CD16, and CD64 on TLR2 gated monocyte subsets.
H) Expression of HLA-DR, CD11c, CD16. and CD64 on DC subsets gated as shown in Figure 2.
I) HLA-DR Expression on monocytes gated by Negative Selection or TLR2 gating.
J) CD163 Expression on monocytes gated by Negative Selection or TLR2 gating.
K) HLA-DR Expression on DC subsets
L) CD163 Expression on DC subsets

We examined the CD11c$^{neg}$ and HLA-DR$^{neg}$ Lin$^{neg}$ population population in the data from Mair et al, [22] but it was not possible to define these cells further.

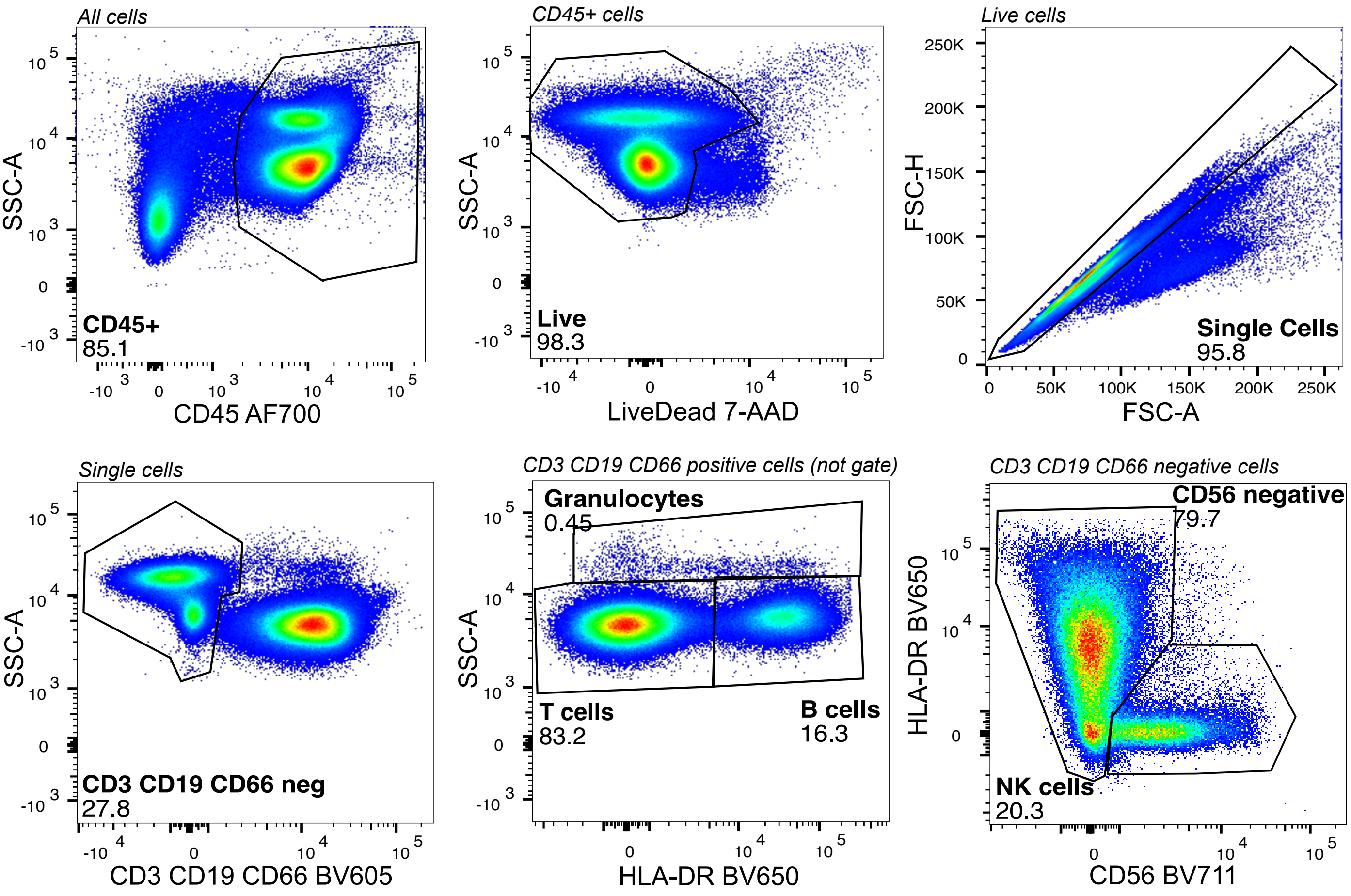


**Supplementary Figure 3:** Gating Strategy for identification of T cells, B cells, Granulocytes, and NK cells.

| Specificity | Purpose | Conjugate | Manufacturer | Clone | Cat | Concentration (µg/mL) |
|---|---|---|---|---|---|---|
| CD141 | mDCs | BV421 | BD Biosciences | 1A4 | 565321 | 0.125 |
| CD14 | Monocytes | V500 | BD Biosciences | MøP9 | 562693 | 3.00 |
| CD3 | T cells | BV605 | BD Biosciences | UCHT1 | 583219 | 0.50 |
| CD19 | B cells | BV605 | BD Biosciences | HIB19 | 740394 | 0.06 |
| CD66 | Granulocytes | BV605 | BD Biosciences | B1.1 | 740423 | 0.04 |
| HLA-DR | Monocytes & DCs | BV650 | BD Biosciences | G46-6 | 564231 | 0.50 |
| CD56 | NK cells | BV711 | BD Biosciences | B159 | 740781 | 0.50 |
| CD16 | Monocytes | FITC | BD Biosciences | 3G8 | 555406 | 1.50 |
| CD1c | mDCs | PerCP-eF710 | Invitrogen | L161 | 46-0015-42 | 0.06 |
| CD163 | Marker of Interest | PE | IQ Products | MAC2-158 | IQP-570R | 0.25 |
| CD11c | mDCs | PE-CF594 | BD Biosciences | B-Ly6 | 564920 | 0.30 |
| CD64 | Monocytes | PE-Cy7 | Biolegend | 22 | B06025 | 0.375 |
| TLR2 | Monocytes | APC | Miltenyi Biotec | REA109 | 130-120-052 | 1.25 |
| CD45 | Leukocytes | AF700 | BD Biosciences | HI30 | 560566 | 0.10 |
| CD123 | pDCs | APC-eF780 | Invtrogen | 6G6 | 47-1239-42 | 0.150 |
| LiveDead | Live cells | 7-AAD | BD Biosciences | | | |

**Supplementary Table 1**: Overview of Flow Cytometry Antibodies
